# The impact of endogenous retroviruses on nuclear organization and gene expression

**DOI:** 10.1101/295329

**Authors:** Ramya Raviram, Pedro P Rocha, Vincent M Luo, Emily Swanzey, Emily R Miraldi, Edward B Chuong, Cédric Feschotte, Richard Bonneau, Jane A Skok

**Author notes:** Contributed equally (Ramya Raviram, Pedro P Rocha). Email Addresses.

## Abstract

**Background:** The organization of chromatin in the nucleus plays an essential role in gene regulation. When considering the mammalian genome it is important to take into account that about half of the DNA is comprised of transposable elements. Given their repetitive nature, reads associated with these elements are generally discarded or randomly distributed among elements of the same type in genome-wide analyses. Thus, it is challenging to identify the activities and properties of individual transposons. As a result, we only have a partial understanding of how transposons contribute to chromatin folding and how they impact gene regulation.

**Results:** Using adapted PCR and Capture-based chromosome conformation capture (3C) approaches, collectively called 4Tran, we take advantage of the repetitive nature of transposons to capture interactions from multiple copies of endogenous retrovirus (ERVs) in the human and mouse genomes. With 4Tran-PCR, reads are selectively mapped to unique regions in the genome. This enables the identification of TE interaction profiles for individual ERV families and integration events specific to particular genomes. With this approach we demonstrate that transposons engage in long-range intra-chromosomal interactions guided by the separation of chromosomes into A and B compartments as well as topologically associated domains (TADs). In contrast to 4Tran-PCR, Capture-4Tran can uniquely identify both ends of an interaction that involve retroviral repeat sequences, providing a powerful tool for uncovering the individual TE insertions that interact with, and potentially regulate target genes.

**Conclusions:** 4Tran provides new insight into the manner in which transposons contribute to chromosome architecture and identifies target genes that transposable elements can potentially control.

## Background

The structural organization of the genome is regulated at different levels to establish a functional framework that facilitates cellular processes such as gene expression and programmed somatic recombination. Fluorescent in-situ hybridization (FISH) studies revealed that chromosomes occupy discrete territories with very little intermingling between them. With the development of chromosome conformation capture (3C) techniques that rely on crosslinking of chromatin in close spatial proximity, additional levels of organization were described. There are many different 3C-based variants, with Hi-C being the most comprehensive, in that it can potentially identify all pair-wise chromatin interactions in a given population of cells [1–5]. The first Hi-C study revealed that each chromosome is divided into active (A) or inactive (B) compartments that range in size from ~5-10MB in mammalian cells. Furthermore, these analyses demonstrate that regions on the chromosome belonging to the same compartment preferentially interact with each other. With improved Hi-C sequencing depth, the presence of topologically associated domains (TADs) were defined. The latter consist of highly self-interacting regions separated from each other by insulated boundaries. Unlike A/B compartments, which are cell type specific, TADs are for the most part invariant across cell-types and orthologous genomic regions of different species. It is thought that the main function of TADs is to restrict the influence of enhancers to genes found in the same domain. Indeed, approximately 90% of promoter-enhancer interactions occur between elements in the same TAD [6–10]. Further support for TAD restricted regulation comes from several studies in which disruption of TAD borders has been shown to lead to aberrant gene expression through exposure to previously insulated enhancers [11–14].

A large portion of a typical mammalian genome is comprised of transposable elements (TEs) however they are typically ignored in high-throughput sequencing-based studies due to their repetitive nature [15, 16]. As a consequence of this, few studies have analyzed how TEs influence, and are influenced by nuclear organization [17, 18]. These mobile elements, which have been propagated in the genomes of all eukaryotic species, can be classified as either DNA transposons or retrotransposons, depending on their mode of transposition. DNA transposons propagate via a cut and paste mechanism while retrotransposons use a copy and paste mode of action that relies on an RNA intermediate [19]. TEs have largely been considered as inert elements that remain silenced, except during a short temporal window during germ cell development. However, this view is changing with increasing evidence demonstrating that not all elements are permanently repressed in the genome. For example, in mammals, transcription and transposition of endogenous retroviruses has been shown to occur at different stages of embryogenesis as well as in adult tissues such as neurons [20–23].

Recent studies have suggested that TEs might act as enhancers capable of influencing expression of endogenous genes [24–30]. This is not surprising, as TEs contain *cis*-regulatory elements such as those found at the 5’ UTR of long interspersed nuclear elements (LINEs) and long terminal repeats (LTRs) of endogenous retroviruses (ERVs) [31–34]. These *cis*-regulatory elements have evolved to control TE transcription but they can also influence expression of adjacent ‘host’ genes [35, 36]. In fact, this idea was first put forward by Barbara McClintock who discovered transposons and described them as ‘controlling elements’ because of their ability to influence the expression of maize genes during development [37, 38]. More recently, genome-wide profiling of transcription factor (TF) binding in human and mouse cells has revealed that TEs contribute a significant proportion of sites [39–42]. In addition, a recent study [43] observed enriched binding of STAT1 in primate-specific ERVs called MER41 after interferon gamma (IFNG) treatment in HeLa and other human cells. STAT1-bound MER41 elements were enriched near innate immunity genes and in particular those that respond to interferon treatment. CRISPR deletion of selected MER41 elements was used to examine their influence on neighboring immune response genes. In each case, deletion of the STAT1-bound MER41 element led to downregulation of the interferon-driven transcriptional response of genes in close proximity. This indicates that MER41 retroviral insertions can act as enhancers which control interferon-inducible expression of neighboring genes. Thus, not all TEs are silenced and inert, and, under particular conditions, they can act as regulatory elements, capable of influencing the expression of surrounding genes. However, only a subset of TEs are predicted to have such effects [44]. Therefore, it is essential to develop techniques that determine the identity of potential target genes that come into contact with these elements in order to ascertain which integration events have potential regulatory function. Follow up targeting experiments can then be used to assess the extent to which they contribute to host gene regulation.

3C-based techniques have become instrumental in the identification of enhancers and the genes they regulate [45–48]. Here we introduce two variants of 3C, tailored to identify different aspects of murine and human transposon-mediated interactions. We first describe 4Tran-PCR, which was used to obtain an interaction-profile for different ERV families in murine cells by mapping all interactions from a specific ERV family to unique sequences within the mouse genome. This approach effectively identified strain-specific, polymorphic insertion sites. In addition, 4Tran-PCR revealed that the interaction profile of mouse ERVs follows a similar pattern to that of host genes, such that if an element is located in an active (A) or inactive (B) compartment, it will preferentially contact other loci within the same compartment, regardless of whether the element itself is decorated with active or inactive marks. Furthermore, 4Tran-PCR demonstrates that TE-mediated interactions are locally restricted to the same TAD.

Capture-4Tran uses biotinylated DNA oligonucleotides (oligos) to capture TEs in conjunction with surrounding uniquely mappable sequences, thereby enabling the identification of the interaction profile of a specific transposon copy, unlike the PCR variant of this technique. This provides a powerful tool for uncovering the individual insertion sites that interact with, and potentially regulate target genes. Here we designed an oligonucleotide that captures interactions involving human MER41 elements and used Capture-4Tran to characterize these interactions. We found that a significant fraction of MER41 interactions occur with genes and involve promoters. Many of these harbor FAIRE-seq peaks, some of which overlap with IFNγ inducible STAT1 binding sites, indicating that these TEs have a potential regulatory role. However, the loops are preformed and can be detected in the absence of STAT1 binding, prior to IFNγ induction, indicating that the contacts are not dependent on the presence of this transcription factor. Collectively, 4Tran provides new insight into the role of TEs in shaping genome organization and regulating cellular processes in mammalian cells.

## Results

### Chromosomal interactions involving TEs can be analyzed using 4Tran-PCR and Capture-4Tran

To investigate different aspects of murine and human retrotransposon-mediated interactions we developed two variants of 4Tran. The first, based on circular chromosome conformation capture (4C-Seq) was called 4Tran-PCR (**Fig. 1**). 4C-Seq captures the frequency with which a bait (or viewpoint) physically contacts other locations across the genome within a cell population. Regions surrounding the bait interact at high frequency due to the polymer nature of chromosomes and the 3D organization of the genome into TADs. As such, 4C-Seq baits are characterized by a single region of strong signal that decays with increasing distance from the bait (**Fig. 2a**). Our strategy to probe TE interaction frequency across the genome consists of using 4C-Seq with primers that hybridize to the repetitive region of transposons. Interactions are mapped to unique genomic sequences, while reads that map to multiple genomic locations, including TEs with low-sequence divergent copies are discarded as we cannot identify which insertion site these emanate from. With this method, several bait-like profiles with a strong accumulation of reads surrounding the integration site of a particular TE type are detected (**Fig. 1** and **2a**). This approach is best suited for analysis of TEs with relatively low copy number where it can be assumed that interactions are captured from the nearest insertion site.

**Fig. 1.**
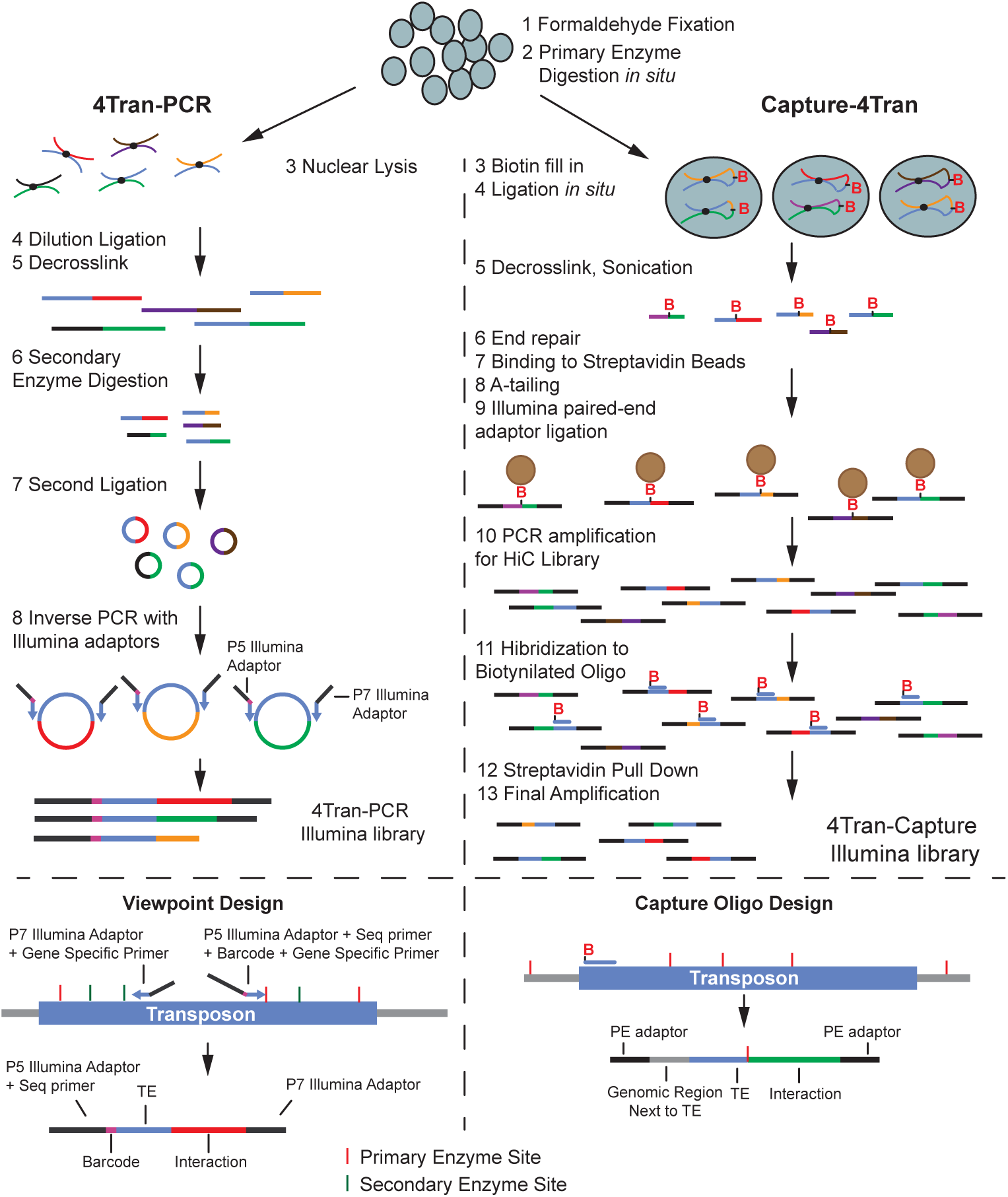
Schematic representation of 4Tran-PCR and 4Tran-Capture approaches. The protocol for 4Tran-PCR is the same as described in [86] for both template preparation and library amplification. Although not shown here, we also successfully tested *in situ* ligation by simply omitting the SDS treatment step following digestion with the primary restriction enzyme. Scheme shows amplification using Illumina single end reads. The protocol also works with paired-end adaptors and for newer Illumina machines these paired-end adaptors are necessary. Capture-4Tran is similar to Hi-C and Capture-C protocols. Our strategy to identify both an interacting fragment and a specific TE integration consists of designing a probe close to the 5’ or 3’ end of the transposon. With this strategy most reads will contain an interaction fragment, part of the transposon to which the oligonucleotide probe hybridizes and the genomic region immediately adjacent to the integration event that can identify one side of an interaction containing the TE.

**Fig. 2.**
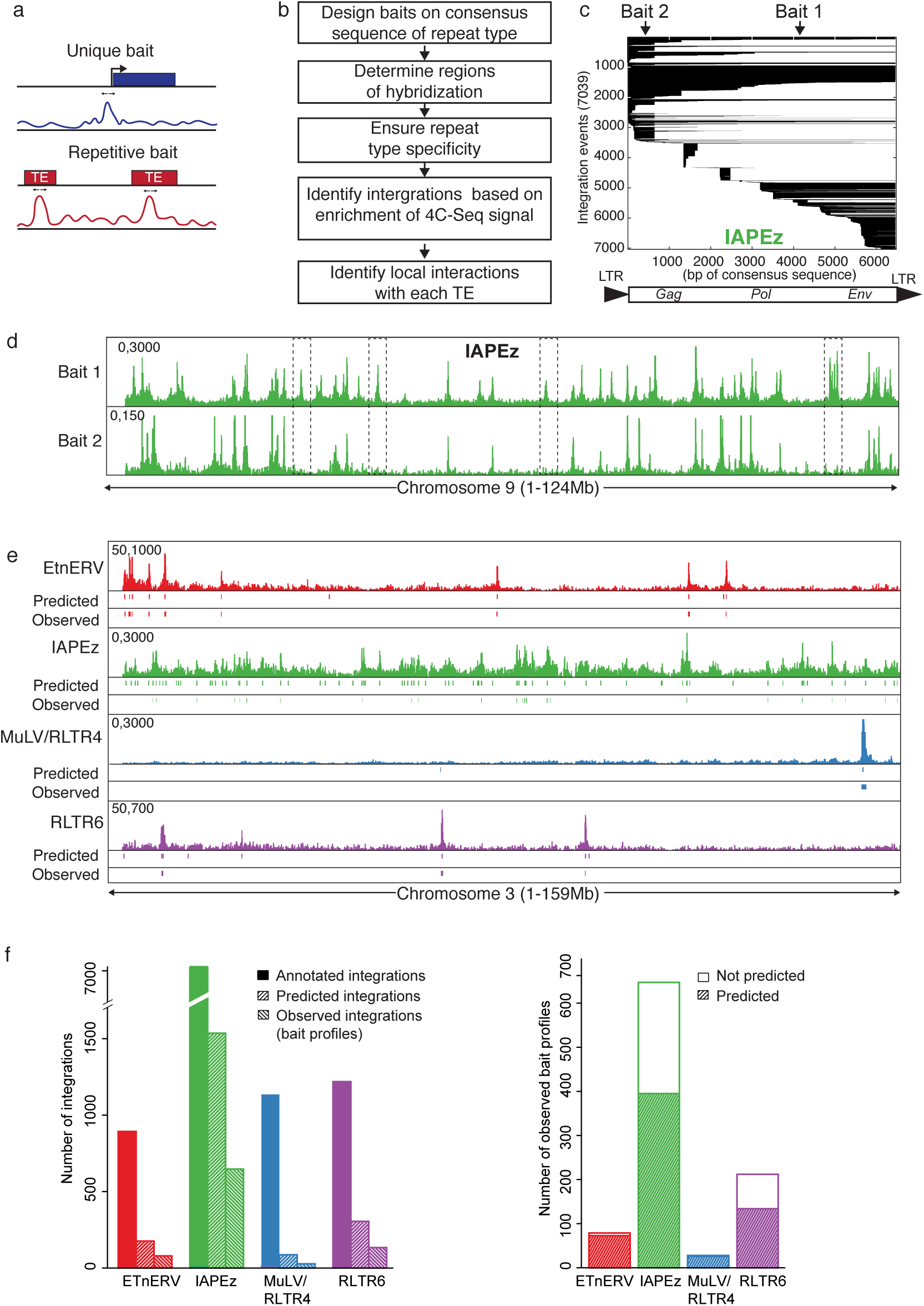
Chromosomal interactions involving TEs can be analyzed using 4Tran-PCR. **a** Scheme showing an accumulation of reads next to where the bait is designed which is typically seen in 4C-seq using a unique bait sequence. 4Tran-PCR results in several bait-like profiles with a large local accumulation of reads in places where there is a TE integration event that corresponds to the primers used. The scheme shows how baits are designed within the transposon as well as the potential location of the two primers and the restriction enzyme sites necessary for amplification of a 4Tran-PCR library. The bottom of the scheme displays an interaction identified by Illumina sequencing containing the barcode, the sequence corresponding to the TE bait and the interaction fragment captured with the bait. **b** Workflow for 4Tran-PCR design and analysis. **c** Schematic representation of all IAPEz-int integration events. Each line represents a different integration and the black lines show which part of the consensus sequence (shown under the plot) is retained by each integration. Integration events are sorted by 5’ position on the consensus sequence and by size of integration. Arrows represent the two locations tested for IAPEz baits. **d** 4Tran data in ES cells for the two IAPEz baits. Boxes represent three integration events detected by bait 1 that are not captured by bait 2. **e** 4Tran data for four different mouse ERVs on chromosome 3. Regions predicted to form bait-like profiles based on the presence of primers sequences (Predicted) and our algorithm based on bait like profiles (Observed) are shown under each 4Tran signal plot. **f** The left plot shows: the number of annotated integrations in the mm10 genome for each ERV (annotated), the number of predicted integrations and the number of observed bait-like profiles. The plot on the right shows the number of observed and predicted bait-like profiles.

As an alternative approach, we developed a 4Tran variant named Capture-4Tran (**Fig. 1**). As the name implies, this is a Capture-C based approach that uses two rounds of DNA capture with 120 bp biotinylated DNA probes to pull down interactions with specific regions of interest [49]. As in Capture-C, our method relies on sonication instead of the second digestion step used in 4C-Seq. This substantially improves our ability to distinguish between PCR duplicates and unique interactions [49]. Additionally, as in Capture Hi-C, we used Hi-C libraries [45] instead of the 3C libraries used in Capture-C. The primary difference between the two approaches is the use of biotin/streptavidin beads in the Hi-C library, which enrich for true ligation junctions formed due to 3D proximity. This increases the number of informative reads obtained at the end of the experiment, which is essential when enriching for interactions from thousands of capture regions. By combining paired-end sequencing with biotinylated oligonucleotides that hybridize to the 5’ or 3’ end of a TE integration site, isolated fragments contain the end of a transposon and the region neighboring the integration site as well as any interacting fragments (**Fig. 1**). This approach increases the chances that uniquely mappable regions are captured with interactions involving TEs and therefore, each interaction can be assigned to a specific TE locus.

### 4Tran-PCR detects annotated ERV integration sites

The workflow for 4Tran-PCR shown in **Fig. 2b** depicts the sequential process from primer design (following 4C template generation) to identification of interactions that we used in 4Tran-PCR. Briefly, we design baits matching the consensus sequence of a TE family, as described in Repbase [50]. We predict which regions of the genome should be amplified with these primers and then intersect these locations with all annotated TEs to ensure that only the TE of interest is captured. Primer pairs that pass all of the above criteria can then be used to detect bait profiles based on enrichment of 4C-Seq signal.

To implement 4Tran-PCR in mouse cells, we focused on TEs classified as endogenous retrovirus (ERVs), which are known to be particularly diverse, recent and active in the murine genome [31]. We selected the following murine ERVs: IAPEz, ETnERV, RLTR6 and MuLV/RLTR4 as these are amongst the youngest murine TE elements and they still retain the ability to transpose. Thus, distinct insertions belonging to the same family are expected to exhibit high sequence similarity, which facilitates repetitive bait design. To analyze their interaction profiles, we designed 4C-like baits for each repeat type, based on the consensus sequence for a set of primary and secondary restriction enzymes. Primer design rules are largely similar to those used in traditional 4C-Seq [51], except that primers are not excluded if they are located in repetitive genomic regions (details in methods section).

Primer pairs were selected based on the following criteria: location within the TE, number of integration copies that can be potentially captured and specificity to the transposon of interest. In addition, we selected baits located close to the 5’ and 3’ ends of TEs where *cis*-regulatory elements recognized by transcription factors tend to occur [52]. It is important to note that due to truncations, the majority of TE insertions are not representative of the full-length element (**Fig. 2c** and **Figure S1a**). For ERVs, these include a high frequency of solitary LTRs, which result from non-allelic recombination events between the LTRs of full-length proviral integrants [31]. Finally, to ensure that no other TE families would be amplified by our bait sequences, we adapted the UCSC in-silico tool to predict bait location and cross-referenced this to the annotation of all murine TEs. We applied these criteria to the baits designed for the four TEs, IAPEz, ETnERV, RLTR6 and MuLV/RLTR4. **Figure 2c** shows the location of each bait relative to the consensus sequence and integration events of all IAPEz elements. The primer location for other ERVs can be found in **Figure S1a**.

To test the primer pairs selected for each ERV, we performed 4C-Seq in mouse embryonic stem cells using DpnII and Csp6 as the primary and secondary restriction enzymes. As predicted, 4Tran-PCR primers located within sequences of endogenous retroviruses generated several bait-like profiles, instead of a single bait profile as in conventional 4C-Seq (**Fig. 2d and Figure S1c**). It should be noted that the different primers for the same transposon did not capture all the same integration events, but rather captured only integration events that retained the primer sequence matching the consensus sequence (**Fig. 2d**). Since only unique reads are mapped, we detect interactions involving the regions surrounding the TE, while those involving only the repetitive regions of the transposon are discarded. **Supplementary Fig. 1b** shows a zoomed in view of an annotated ERV bait showing the 4Tran-PCR signal derived from unique sequence reads of surrounding regions and an absence of TE reads in the center.

To determine if 4Tran PCR can identify annotated ERV integration events we selected preferred primer pairs based on their proximity to LTRs and their ability to generate efficient 4C signal amplification. As the mm10 reference genome from the Genome Reference Consortium is based on the C57/Bl6 strain we performed 4Tran-PCR using template prepared from murine *ex vivo* derived splenic resting B cells from mice of this strain. B cells were chosen for their easy availability in our lab and because high-resolution Hi-C datasets that we could use as validations were restricted to B cells at the time of these experiments. Our first goal was to test whether the observed 4Tran bait profiles align with known ERV integration sites (**Fig. 2e**). To identify these regions in an unbiased manner, we binned the genome into 200kb windows and compared the 4Tran signal in each window to a background distribution. Windows that contained enriched signal were called “observed baits”. Additionally, we defined “predicted baits” as ERV integration events that retain the region of the consensus sequence matching the bait primers (see methods section for details).

The ERV family with the highest number of predicted integration events is IAPEz, followed by RLTR6, ETnERV and MuLV. The number of observed baits for each of these ERVs also followed the same trend (**Fig. 2f left**). For all four ERVs analyzed we detected fewer observed baits than predicted. This could either be due to emergence of mutations that are not annotated in the reference genome, or to polymorphisms in our C57/Bl6 colony. Another possibility is that our method may not distinguish some of the signals as true 4Tran bait profiles due to low signal. This is particularly relevant for samples with a high frequency of integration sites where the higher density of bait-like profiles could obscure the distinction between bait signal and regions of high interactions. Another potential source for the disparity between the number of “observed” and “predicted” baits is that multiple integrations in close proximity cannot be detected as single copies.

Finally, we asked whether the locations we identified as observed baits had been predicted. (**Fig. 2f** right). The ERV elements with fewer observed baits, ETnERV and MuLV/RLTR4 displayed an almost perfect correspondence between observed and predicted baits. In contrast, the RLTR6 and IAPEz ERVs, had a higher number of observed baits, many of which were not predicted. As mentioned above, this discrepancy could be due to the high density of bait-like profiles, which blur the distinction between bait signal and regions of high interactions. Thus, the use of 4Tran is best suited to ERVs with a lower number of integration sites. Alternatively, primers that detect fewer integration sites of a specific TE family are preferred to primers that detect all hybridization sites.

### 4Tran-PCR detects TE insertion polymorphisms

As 4Tran is able to detect the location of ERV integration events through enrichment of their local chromatin interactions we next asked whether differences in integration sites of mobile ERVs could be detected. Since the four ERV elements we analyzed are among the youngest in the murine lineage we tested this hypothesis by performing 4Tran in *ex-vivo* derived resting splenic B cells from the C57/Bl6 and 129S6 mouse strains. For this, we used the same primer sets described above to analyze integration of ETnERV and MuLV/RLTR4. We found many observed baits that were shared between mouse strains as well as C57/Bl6 and 129-specific integrations (**Fig. 3a**). As expected, the C57/Bl6 strain on which the mm10 reference genome is built performed better in identifying observed versus predicted baits compared to the 129S6 strain (**Fig. 3b**).

**Fig. 3.**
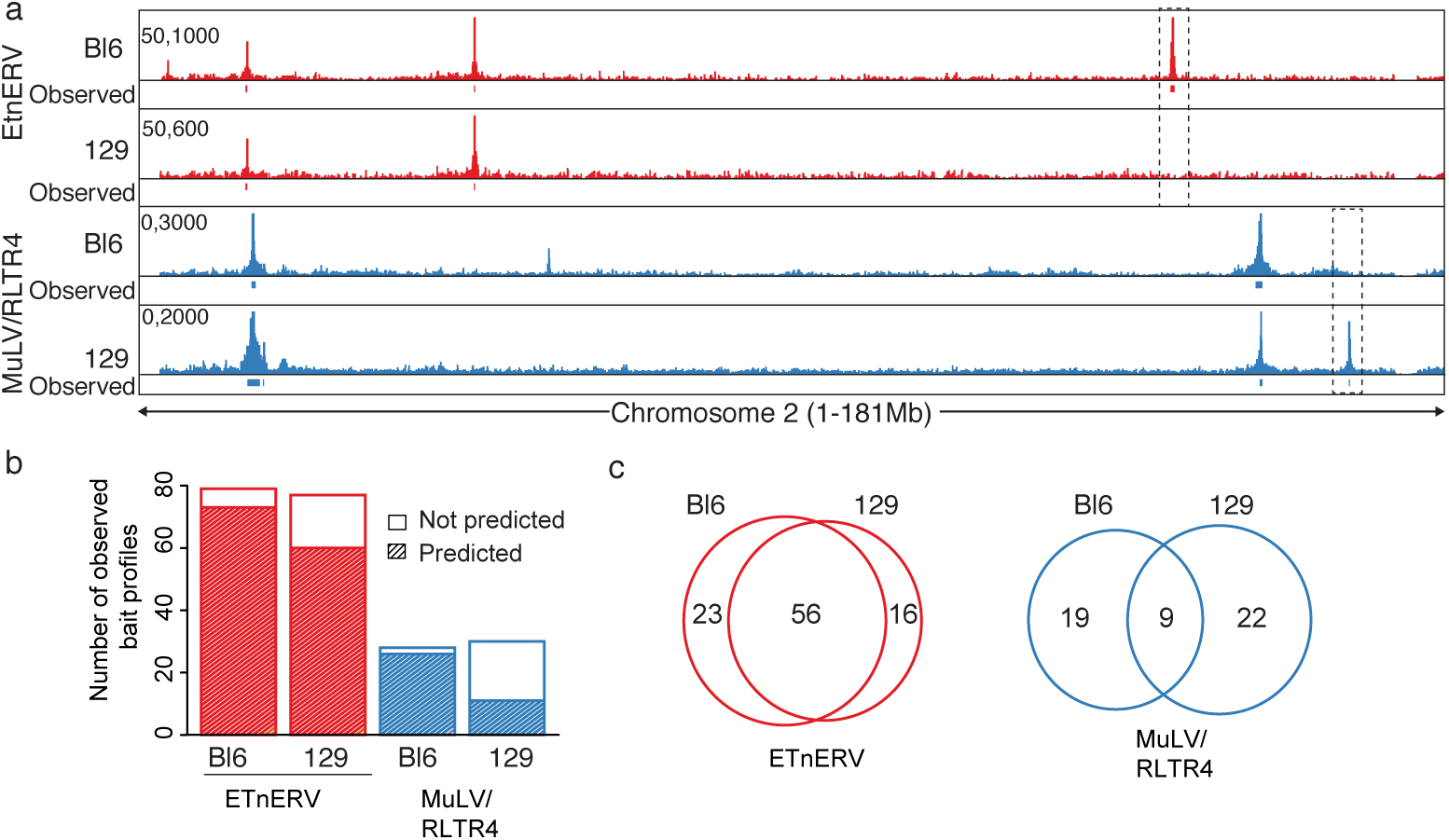
*4Tran-PCR detects TE insertion polymorphisms* **a** Raw 4Tran reads are shown for the same baits (ETnERV and MuLV/RLTR4) in splenic B cells isolated from mice of either the Bl6 or 129 strains. Boxes represent regions where either Bl6 or 129-specific integrations were detected. **b** For these 4 datasets we show how many of the observed integrations were predicted based on their sequence. **C** Venn diagrams depicting how many observed bait-like profiles are shared between the Bl6 and 129 strains for each of the two ERVs.

The Mouse Genomes Project (sanger.ac.uk/science/data/mouse-genomes-project) database, which generated whole-genome sequencing of 18 different mouse lab strains [53] was used to confirm the presence of structural variations in non-overlapping regions. For example, in the top panel of **Figure 3a**, an ETnERV integration is detected only in the C57/Bl6 strain. When this bait location was compared between mouse strains in the database, we found a deletion that coincides precisely with the C57/Bl6 annotated ETnERV integration in the 129 mouse strain. Conversely, in the bottom panel of **Fig. 4a** we show an MuLV/RLTR4 bait detected in 129S6 but not in C57/Bl6 cells, consistent with the annotation of a polymorphic insertion at this location in the database. These results indicate that differences in observed baits across strains correspond to strain-specific ERV insertions.

**Fig. 4.**
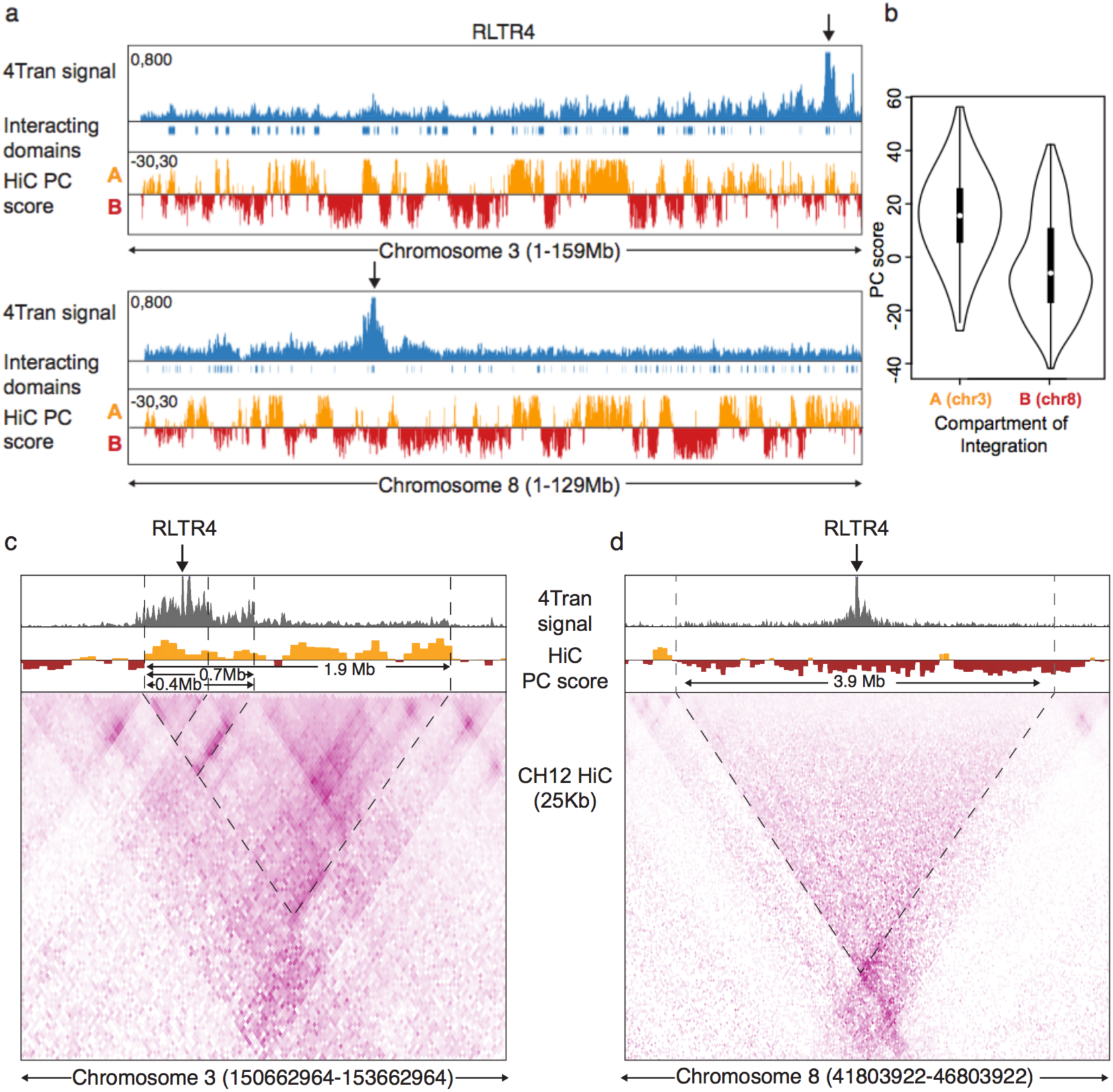
ERV interactions are constrained by the different levels of nuclear organization **a** Whole chromosome view of 4Tran-PCR signal for RLTR4 integrations on chromosomes 3 and 8. The single integrations shown for these chromosomes are highlighted with an arrow over the plot and the regions identified as significantly interacting with these sites are shown under 4Tran-PCR signal as boxes. Hi-C data is represented by the PC-score calculated for each 50kb bin. A positive PC score is characteristic of A regions, while a negative score is associate with B regions. **b** Violin plots representing the PC score for all regions identified as interacting *in cis* with the RLTR4 integration in chromosomes 3 and 8. An integration in compartment A leads to contacts with other compartment A regions, while the reverse is true for an integration in compartment B on chromosome 8. **c** and **d** High resolution 4Tran-PCR data is shown together with Hi-C from Ch12 B cells. Hi-C is shown using both principal component score and 25 Kb-bins. Dashed lines highlight regions of high 4Tran-PCR signal and borders of domains as described in the results section.

When differences between the strains were quantified genome-wide, ETnERV showed a higher level of conservation between 129 and C57/Bl6 than MuLV/RLTR4. As shown in **Fig. 3c**, up to 70% of the MuLV integrations were strain-specific, in contrast to the ETnERV family where 70% of integration sites are shared between the two strains. These frequencies are in line with previous estimates for each of these ERV families of the numbers of elements that have polymorphisms across different mouse sub-strains [54].

Finally, we tested 4Tran-PCR in different sub-strains of 129 mice using the ETnERV bait, and found that polymorphic insertions could be detected between sub-strains. In the left panel of **Figure S2a**, an integration event is captured in 2 out of 3 129 sub-strains while in the right panel, another integration is captured in only 1 out of 3. We cross-referenced the regions of differential integrations to the Mouse Genomes Project database and found that the annotations matched our findings. Surprisingly, we also detected differences in integration events between littermates using the MuLV/RLTR4 bait and an integration on chromosome 2 was identified in one mouse that was not detected in its littermate (**Figure S2b).** This transposition event is not annotated in the reference C57/Bl6 genome and likely arose in the germ cells of one of the parents. In sum, these examples show that 4Tran-PCR is capable of detecting differences in TE integration sites that are present in a population of cells. While other techniques have been developed solely focused on detecting new TE integrations [55, 56], a more comparative analysis is needed to assess the performance of this aspect of 4Tran-PCR.

### ERV interactions are constrained by the compartments and local domain structure

Having determined that 4Tran-PCR identifies chromosomal interactions of ERV elements we looked at how transposon contacts are influenced by the different levels of nuclear organization and asked whether ERV-mediated long-range interactions are influenced by chromosome compartmentalization. Compartment formation (A and B) seems to be mostly guided by the propensity of large chromatin fragments with similar histone and DNA modification patterns to share the same physical space. Formation of compartment A, for example, results from the spatial clustering of genomic regions enriched for histone post-translational modifications associated with transcriptional activity, such as H3K4me1/3 or H3K27ac [5, 6]. Therefore, we asked whether full-length ERV copies known to be decorated with the heterochromatin mark H3K9me3 [52, 57] but located in A compartment regions would frequently contact regions in the B compartment that share H3K9me3 enrichment, or rather if they would evade compartmentalization because of the influence of their local A compartment neighborhood.

To address this question we used Hi-C and ChIP-seq data from a splenic murine lymphoma IgM+ B cell line, CH12 that is used as a model for B cells in the same developmental stage as those we use here [58]. We performed a principal component analysis on 200 Kb-binned Hi-C data and classified each bin as compartment A and B depending on a positive or negative principal component score, respectively [59] (**Fig. 4a**). We then focused on data generated using the RLTR4 bait where our primers detected a single integration site on chromosome 3 that is located in a region that belongs to compartment A (**Fig. 4a**). Full-length insertions of this ERV are silenced in B cells and decorated with the heterochromatin mark H3K9me3 (**Figure S3**). Visual inspection of the interactions of this RLTR4 copy with the rest of the genome, revealed that the regions of higher interaction belonged to the A compartment. The 4C-ker pipeline [60] was used to quantify this and determine which regions on chromosome 3 interact at high frequency with the MuLV integration site. As shown in **Fig. 4b**, interacting regions have positive PC score values, indicating that this ERV integration preferentially interacts with other compartment A regions on chromosome 3. This data demonstrates that, just like non-repetitive loci, TE long-range interactions are guided by their compartment status and determined by the overall epigenetic status of their compartment rather than their specific chromatin marks. As a validation, interactions of an RLTR4 copy integrated on chromosome 8 in a B compartment region were examined. Again, these reveal that most interactions occurred with regions in the same compartment (**Fig. 4a** and **Fig. 4b**). A comparable analysis with two RLTR4 integrations on different chromosomes further validated the finding that ERV long-range interactions are restricted to regions within the same A or B compartment that they are integrated in, rather than to regions with the same epigenetic state.

We then asked whether the local interactions of transposable elements are confined within smaller architectural structures such as TADs [61, 62]. Here we took advantage of the high resolution Hi-C CH12 data to directly compare with 4Tran-PCR signal. Hi-C data revealed that the RLTR4 integration on chromosome 3 element is integrated in a small domain of approximately 0.4 Mb which is insulated from its immediate upstream region and is a substructure of two wider (0.7 and 1.9 Mb) nested domains that expand downstream of the MuLV integration site (**Fig. 4C**). 4Tran-PCR signal portrays the same architecture, with the strongest signal located within the 0.4 Mb domain. Interactions are dramatically reduced in the regions immediately upstream, indicative of strong insulation of the 0.4 Mb domain from its flanking regions. Furthermore, even though the regions upstream of the RLTR4 integration are closer on the linear chromosome, 4Tran signal is much stronger downstream of the RLTR4 element and constrained by the two subdomains of the larger 1.9 Mb domain. In contrast, the RLTR4 integration on chromosome 8 is much less structured, likely because it is located in a large 3.9 Mb compartment B domain (**Fig. 4D**). Similarly, 4Tran signal from this RLTR4 integration decays linearly and symmetrically with distance from the ERV site and is constrained by the borders of the domain detected by Hi-C. Taken together these results indicate that both local domain structure and organization of the genome into compartments influences the manner in which TEs interact with other loci. It is important to note that low sequencing depth 4Tran signal (2-10 Million reads) achieves a similar resolution to billions of Hi-C reads. Thus, 4Tran-PCR can be used to easily assess both local and long-range interactions from transposable elements.

Identifying interactions from individual TEs using 4Tran-PCR works optimally if insertion sites are well separated on the linear chromosome. However, 4Tran-PCR can confound the characterization of interactions from TE integrations in close proximity and it is thus best suited for analysis of TEs with relatively low copy number integrations. Under these circumstances it can be assumed that interactions are captured from the nearest site as by definition, regions in the same domain interact at higher frequency than with the rest of the genome. A further drawback of 4Tran-PCR is that it is less efficient at amplifying interactions from families with highly divergent sequences, as it relies on the presence of two PCR primers and two restriction enzyme sites. To address these issues we developed a variant of 4-Tran, Capture-4Tran.

### Capture-4Tran identifies pairwise interactions from TEs

Capture-4Tran overcomes the issue related to repetitive sequence mappability by combining paired-end sequencing with biotinylated oligonucleotides that hybridize to the 5’ or 3’ end of a TE integration site. It can therefore be used to uniquely identify interactions from individual TE insertions. To test the feasibility of this approach, we first performed Capture-4Tran for the IAPEz family, which is also one of the youngest and more homogeneous ERVs in the mouse genome. We performed a pilot experiment at low-sequencing depth using a probe based on the consensus sequence of the LTR that is most commonly associated with IAPEz elements, IAPLTR1a. As shown in **Figure S4,** this strategy allowed unique mapping of interactions from IAPLTR1a elements, consisting of either a solo LTR or full-length insertions. Up to 30.5% of all interactions that we uniquely aligned in our Capture-4Tran experiment have at least one side mapped to an IAPLTR element demonstrating the ability of our probe to enrich for the desired fragments.

We next tested if Capture-4Tran is able to identify interactions from slightly older TEs that are therefore less homogeneous in sequence, but may have greater potential for being coopted for adjacent gene regulation [34, 40, 63]. For this we focused on the MER41 LTR elements in the human genome. MER41 represents the LTR of an endogenized gammaretrovirus that entered the anthropoid primate lineage 45-60 million years ago [43]. In the human genome there are currently 7190 MER41 copies divided into 6 subfamilies (A-E, G), which range on average from 83-91% nucleotide similarity to their respective consensus sequence [50]. To identify MER41-mediated interactions we designed a probe of 120 nucleotides around Dpn2 restriction enzyme sites in the MER41B consensus sequence (a subfamily of MER41 that contains STAT1 binding sites). HeLa cells treated with Interferon gamma (IFNγ) were selected to test the ability of this probe to enrich for MER41 interactions as this system was previously used to demonstrate the involvement of a subset of MER41 elements as interferon-inducible enhancers regulating adjacent immune genes [43]. Using Blastn, we predicted that 3452 (49.8%) individual MER41 copies would be captured by our probes (1790 MER41A, 1609 MER41B, and 180 MER41C) (**Fig. 5b**).

**Fig. 5.**
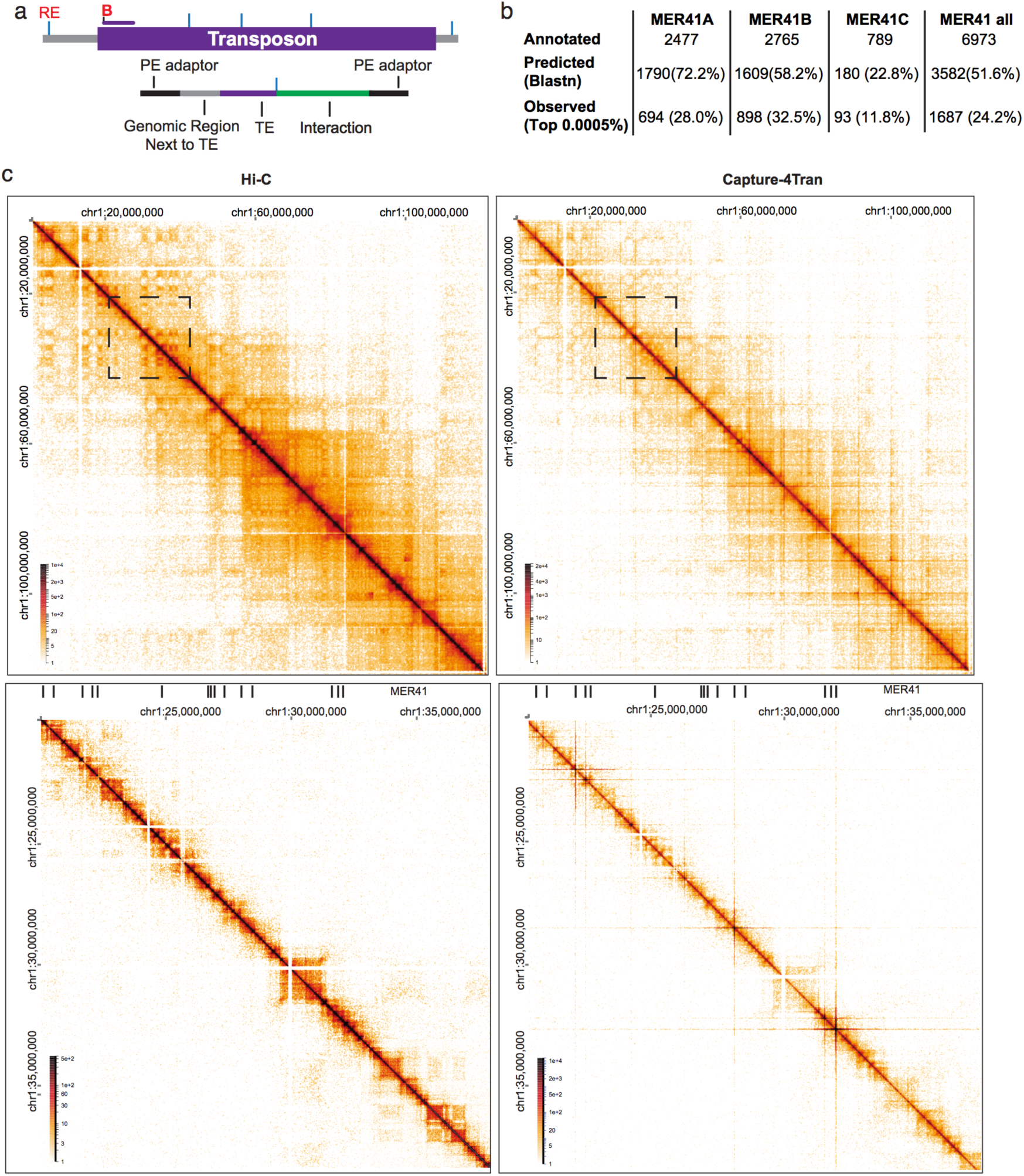
Capture-4Tran identifies pairwise interactions from TEs **a** Scheme of probe design for Capture 4Tran. Blue lines represent the location of the restriction enzyme site. Biotinylated oligonucleotide are shown with a letter B. This procedure generates reads associated with the TE of interest, the genomic region adjacent to the TE, and an interaction involving the TE. **b** Table representing number of annotated, predicted and observed MER41 LTR elements. The 3 MER41 subfamilies that our MER41 oligonucleotide binds to is shown. Percentages relate to the total number of annotated MER41 elements in HeLa cells. **C** Hi-C and Capture-4Tran using the MER41 probe. Boxes in the top panels show zoomed in views from the bottom panels. Data from two replicates was combined for visualization.

In addition to the Capture-4Tran library, we sequenced a Hi-C library for comparison. Capture-4Tran yielded 25 times more sequencing reads associated with MER41B-mediated interactions than the Hi-C approach (5 million unique contacts out of 19.0 million versus 55.0 thousand out of 61 million contacts in Hi-C). The reads generated a contact matrix that is clearly distinct from a Hi-C matrix in that TADs and compartments are no longer distinctly visible. Instead, strong horizontal and vertical lines stand out that are centered at the diagonal of the matrix and emanate from MER41 elements (**Fig. 5c**). To determine whether the MER41 probe hybridized to Dpn2 fragments not predicted by Blastn analysis, we calculated the number of reads per Dpn2 fragment and considered only those belonging to the top 0.005% (see method for details). Altogether 1766 Dpn2 fragments representing 1687 distinct MER41 LTR elements were detected. In addition, 272 fragments that were not predicted by Blastn were identified, of which 183 contained MER41 elements and the remaining 86 (representing less than 5% of all captured fragments) corresponded to regions without annotated MER41 elements (**Fig. 5b**). Thus, using a single oligonucleotide we are able to detect a substantial fraction of copies (24.2%) from an abundant and relatively ancient TE family with a high rate of specificity (95%). The small percentage of off-target enrichment suggests that inclusion of a few extra probes hybridizing to other regions of the consensus sequence, or targeting the consensus sequence of other MER41 subtypes (here an oligo targeting only MER41A B and C was used) would easily allow capture of the remaining family of MER41 LTR elements.

### Characterization of significant interactions from MER41 elements

The Chicago pipeline was used to identify specific pairwise interactions between MER41 elements and other genomic regions [64]. Using a Chicago score threshold of 7.5, 4943 interactions from 944 MER41 elements with a mean value of 5.2 interactions (median value of 2) per bait were identified (**Figure S5a**). The majority of capture-baits occurred on MER41B and MER41A elements (~97%), with a few on MER41C, MER41D and MER41-int (**Fig. 6A**). MER41 capture-bait positions were overlapped with STAT1 ChIP-Seq and FAIRE-Seq data (a proxy for accessible chromatin regions) from HeLa cells treated with IFNγ for 4 hrs (data obtained from ENCODE). In line with recent reports that suggest a regulatory role for MER41 elements, 47% of capture-baits were found to overlap with either FAIRE-Seq or STAT1 signals. MER41B elements, which contain a STAT1 binding motif and have been shown to preferentially bind STAT1, showed greater overlap with STAT1 ChIP-Seq peaks, than MER41A elements, which lack the STAT1 binding motif (**Fig. 6A**). As expected, ~75% of interactions occurred between loci within the same TAD (**Fig. 6B**). In line with this observation, the median distance between MER41 capture-baits and their *cis* interactions was found to be ~162.6 kb (**Fig. 6C**). Next, we asked whether loci that interact significantly with MER41 are significantly enriched for STAT1 binding and open chromatin marks. The overlap of interactions with STAT1 peaks, FAIRE-Seq and gene bodies was found to be significant for all these features compared to randomized backgrounds (**Fig. 6D**). This result is in line with previous studies showing that loci bound by the same transcription factor tend to interact at higher frequency. MER41 elements also interacted more frequently with gene-rich regions than expected by chance (**Figure S5B**). Interestingly, 32 interactions were detected between MER41 elements and preferentially occurring within the same sub-family (**Fig. 6E**). An example of this can be seen in the *SERPINB6* gene. A MER41B element downstream of the gene interacts with the promoter of SERPINB6, which contains a truncated MER41B element that is ~60bps long (**Fig. 6F**). Together, these data suggest that MER41 elements can engage in long-range genomic interactions with active regions based on open chromatin and transcription factor binding.

**Fig. 6.**
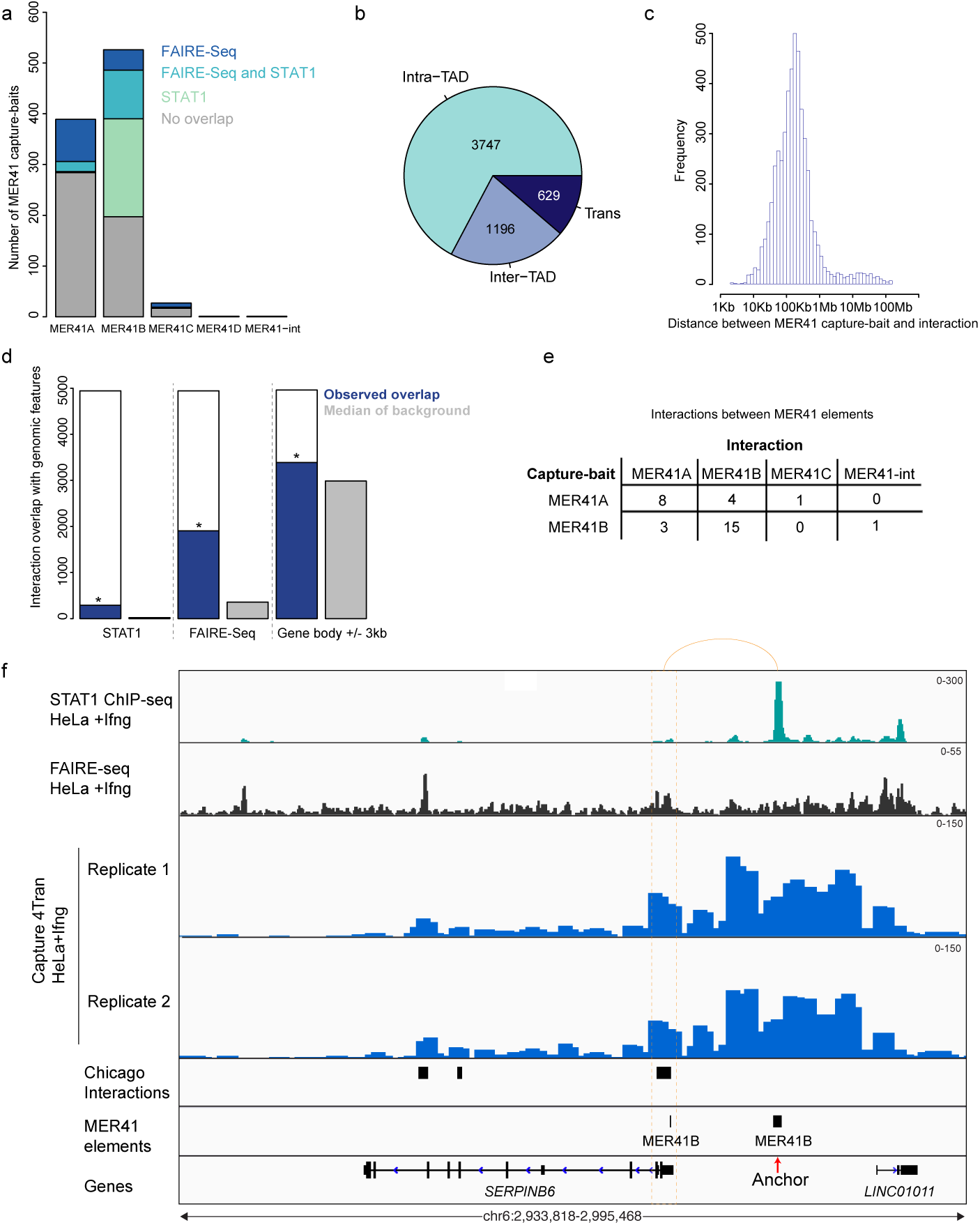
Interactions from MER41 elements. **a** Overlap of MER41 anchors (+/-1kb) with FAIRE-Seq and STAT1 ChIP-Seq peaks. **b** Interactions are classified whether they are in *cis* (same chromosome) or trans (other chromosomes) and *cis* interactions are further divided based on whether they are in the same TAD as the MER41 anchor element. **c** Histogram of the distance between MER41 anchors and their *cis* interactions. **d** Overlap of interactions (+/-1kb) with STAT1 ChIP-Seq peaks, FAIRE-Seq peaks and gene bodies +/-3kb. The background regions were generated based on randomly shuffling the position of the interactions on the same chromosome and calculated the overlap with each feature (median of 1000 iterations displayed). Significance was calculated based on the number of times an overlap with the randomized interaction positions is greater than observed and divided by 1000. **e** Number of interactions from MER41A and MER41B anchors that overlap with any MER41 element. **f** Genomic tracks of STAT1 ChIP-Seq signal, FAIRE-Seq signal, Capture 4Tran data from HeLa cells treated with IFNγ. Red arrow represents the MER41B anchor and the orange dotted rectangle represents the interaction at the *SERPINB6* promoter.

### MER41 elements are involved in several potential regulatory interactions

To determine whether some of these interactions could have regulatory potential, we considered the top 3 ranked *cis*-interactions for each MER41 element with a minimum Chicago score of 10 (default threshold of significance is 5 [64]). In addition, we focused on interactions between MER41 elements with regions mapping within 1 kb of gene promoters. With these more stringent criteria we identified 151 interactions implicating 105 MER41 copies contacting a total of 129 TSSs (**Table S3**). The majority of the MER41 elements and gene promoters overlapped with STAT1 or FAIRE-Seq peaks (**Fig. 7a**). However, only 39 of the 113 expressed genes contained STAT1 or FAIRE-Seq peaks and only 2 of these genes are up-regulated by IFNγ treatment (**Fig. 7a**). This finding suggests that these interactions could be mediated by other TFs or other architectural proteins. It is of note that of the 151 interactions with a potential regulatory role, 113 involve genes expressed in HeLa cells and 74 originate from MER41 elements with either a STAT1 or a FAIRE-seq peak. In general, a 1:1 ratio of MER41 element to promoter interactions was observed, however some elements contact 2 or more promoters (**Fig. 7b**). In addition, most promoters were in contact with a single MER41 element, with only 6 out of 129 contacting multiple MER41 elements (**Fig. 7b**).

**Fig. 7.**
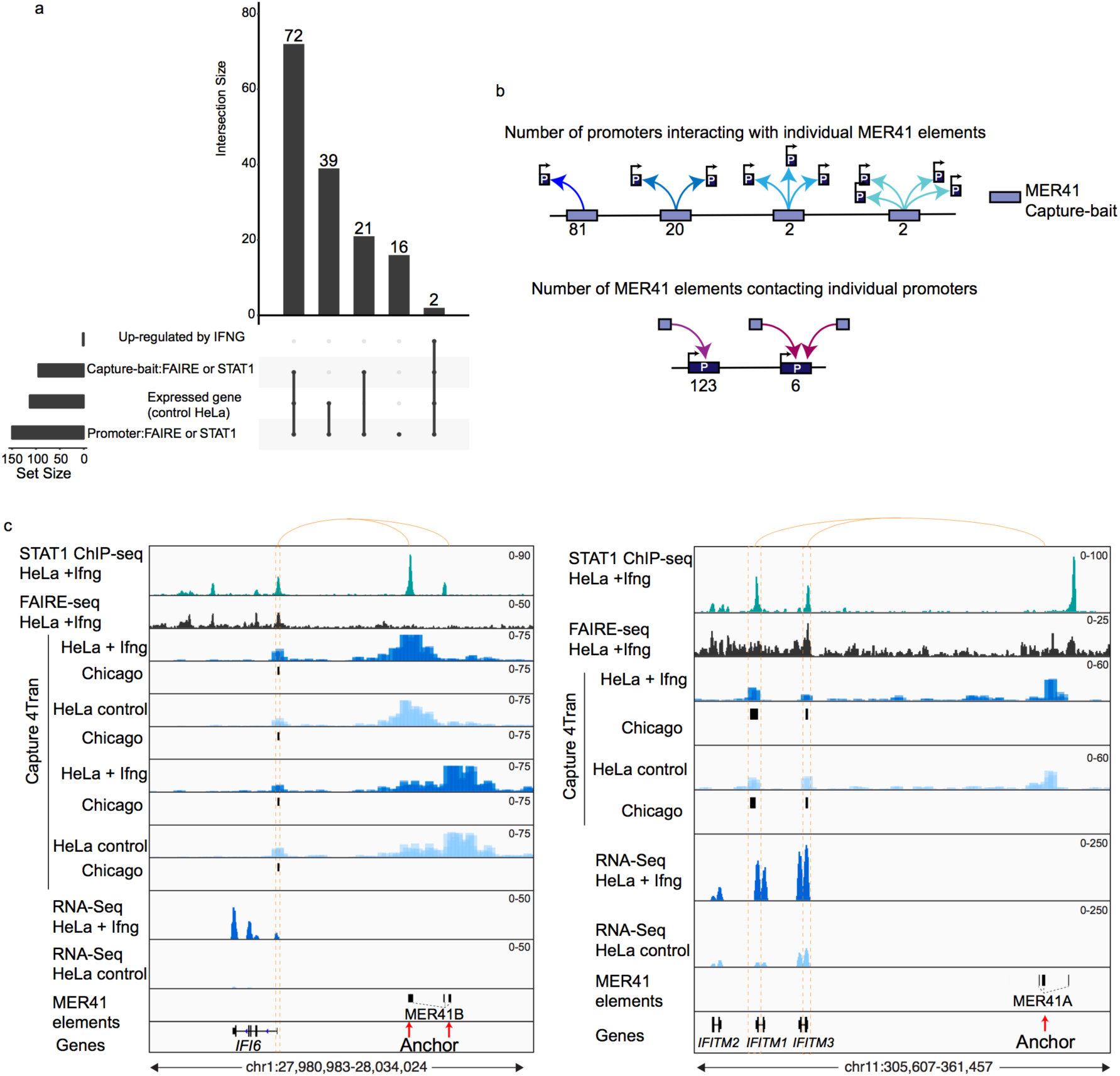
Interactions between MER41 elements and promoters. **a** UpSet plot displaying the overlap between the regulatory interactions with several features. Overlap was calculated for the MER41 anchor side and promoters separately. **b** Breakdown of the number of promoters each MER41 element is interacting with (top panel) and the number of MER41 elements contacting each promoter (bottom pane). **c** Genomic tracks of STAT1 ChIP-Seq signal, FAIRE-Seq signal, Capture 4Tran data from HeLa cells without and without IFNγ treatment. Red arrow represents the MER41B anchor and the orange dotted rectangle represents the interaction at the *IFI6, IFITM1* and *IFITM3* promoter.

Of the 175 genes induced upon interferon gamma (IFNγ) treatment identified by microarray analysis (see methods for details) [65], two genes, *IFI6*, *IFITM1* that robustly respond to IFNγ and significantly interact with a MER41 LTR element were identified (**Fig. 7c**). It has previously been established that a MER41B element located ~20 kb downstream of *IFI6* contributes to its IFNγ -mediated transcriptional upregulation. Specifically, deletion of this element leads to an approximately two-fold reduction in the level of IFI6 IFNγ-mediated transcripts [43]. Capture-4Tran data clearly indicates that this MER41B LTR integration is engaged in an interaction with the promoter of *IFI6*. Moreover, 4Tran identified a second MER41B element inserted further upstream in contact with the *IFI6* promoter, suggesting that this element could also contribute to the regulation of *IFI6*.

4Tran also captured an interaction between the *IFIMT1* and *IFITM3* genes and a MER41A element downstream of these genes (**Fig. 7c**). This element is within a cluster of several MER41 elements and is immediately upstream of a STAT1 binding site. Notably, a previous study aimed at identifying enhancers of the *IFITM* genes, used CRISPR to delete sequences in this region and found sequences overlapping the STAT1 binding site to be an important enhancer of the IFITM1 cluster of genes [65]. Thus, it is possible that the interaction between the MER41A and the IFITM genes is responsible for, or contributes to the contacts between the STAT1 binding region and the IFITM promoters. However, although both *IFI6* and *IFITM1* are strongly upregulated by IFNγ in HeLa cells, the contacts between MER41 elements and the promoters are established prior to treatment, indicating that STAT1 binding is not responsible for initiating the physical regulatory interactions at these loci. Furthermore, differential analysis of all 4943 Chicago-detected interactions using DESEq2 revealed only 6 MER41 anchors with reduced contact frequency after IFNγ induction (see methods for details). One of these involved a promoter for the *FES* gene, which does not respond to IFNγ treatment.

A subset of genes known to be regulated by MER41 elements in response to IFNγ in HeLa cells such as AIM2 [43] were not identified by Capture-4Tran. The most likely explanation is that these elements are too close to their target promoters on the linear chromosome (<5 Kb) to be detected by this assay. Furthermore, some of these genes were shown to be regulated by elements of MER41 subfamilies that our probe does not hybridize to (e.g., the element regulating AIM2 belongs to the family MER41E, which was not targeted by our probe). In sum, our analyses suggests that multiple MER41 elements in the genome might function as regulatory elements, and that Capture-4Tran can be used to identify these elements as well as a list of target genes that could be further validated using other experimental approaches.

## Discussion

We developed a set of 3C-based tools that takes advantage of the repetitive nature of transposons to study how they impact nuclear organization. Our findings provide insight into how transposons interact with other components of the genome, including host genes, and are themselves influenced by chromosome organization. Specifically, we established two different iterations of chromosome conformation capture, a PCR and a Capture-based variant that we collectively call 4Tran. We applied these tools to endogenous retroviruses that have retained the ability to replicate and insert themselves elsewhere in the genome and show that 4Tran can detect both fixed insertions and those that remain polymorphic in the population. Interestingly, young ERVs that have integrated into an active A compartment engage in long-range interactions with other A compartment loci of the genome, despite being heterochromatically silenced (judged by H3K9me3 enrichment). In addition, we find that chromatin interactions of ERVs are mostly delimited by the organization of chromosomes into smaller, highly self-interacting domains and their chromosomal looping is largely restricted to these regions. Finally, we demonstrate that Capture-4Tran can be used to identify TEs that establish direct looping interaction with adjacent host gene promoters and thereby likely modulate host gene transcription.

The choice between use of the PCR-based and the capture-based variants of 4Tran will depend on the biological question. 4Tran-PCR works best with younger TE families that have a moderate number (up to a few hundred) of integration events with high sequence similarity between different copies. It provides high-depth sequencing data at low cost, is easier to implement than Capture-4Tran and can be used to characterize long-range TE contacts and boundaries of domains that restrict their local interaction. Capture-4Tran on the other hand, can be used for analysis of younger TEs, as well as older families with high sequence divergence. Indeed, it bypasses most of the limitations of 4Tran-PCR, including small size and it is highly specific for the elements it captures. As a result, it can be used to study high copy number transposons such as SINEs and LINEs.

A number of recent studies have focused on the role of transposons in gene regulation. In the absence of a chromosome conformation capture tool, TE influence has been examined in the limited context of the nearest gene. However, TEs can participate in both long-and short-range interactions and could potentially be involved in the regulation of multiple target loci. 4Tran therefore provides a tool for more precisely defining the extent of this control and identifying TE-mediated interactions with specific promoter regions.

Capture-4Tran has a clear advantage over promoter Capture Hi-C [45], which enriches for all DNA interactions involving gene promoters. For example, it will only enrich for promoter interactions if a gene is in contact with the TE of interest. This results in reduced numbers of uninformative reads and thus a lower sequencing depth is required, making it more affordable to analyze replicates and compare different conditions. Moreover, by designing a capture probe next to the TE integration site it is possible to detect exactly which repetitive element is engaged in an interaction with a gene promoter, while analysis of promoter-based contacts is less likely to identify interactions with the repetitive portion of younger TEs with low level of sequence divergence between copies.

4Tran results are consistent with the previous proposal that a subset of MER41 elements act as interferon-inducible enhancers that contribute to STAT1-mediated transcriptional activation of interferon sensitive genes in response to pathogen infection. Our results show that MER41 elements function as a prototypical enhancer by making contact with the promoter of flanking genes. Furthermore, our analysis uncovers many direct MER41-gene associations that further support the notion that these primate-specific endogenous retroviruses have a substantial impact on gene regulation in human cells and represent an important force driving the regulatory evolution of the primate innate immune response. These experiments add to a growing body of evidence demonstrating two types of enhancer-promoter contacts: stable and dynamic. Stable contacts such as the majority of enhancer-promoter contacts identified during a TNFα response are formed prior to signaling [66], are in line with our observations for Mer41 in IFNγ stimulated cells. Similarly, genes activated under conditions of hypoxia by the HIF transcription factor are engaged in stable DNA contacts with their enhancer elements [67]. In contrast, DNA enhancer promoter loops can be dynamically rewired during differentiation and cell-type specific DNA contacts are established only when enhancers are activated by binding of cell-type specific transcription factors [62, 68-73]. The situation in *Drosophila* contrasts with the findings from mammalian studies as in early development pre-established loops connect enhancers and promoters even before gene activation [74]. The fact that both interferon and TNFα responsive genes establish loops with similar dynamics suggests that there is a fundamental difference between transcriptional responses to stress/pathogens and those underlying developmental transitions. It is tempting to speculate that the interferon response is response is similar to TNF-α signaling and that there is a fundamental difference between transcriptional responses to stress/pathogens and those underlying developmental transitions.

Despite major advances in the field since the development of chromosome conformation, there is still a lot to be learned about the role of nuclear architecture in gene regulation. In particular, our knowledge of the contribution of TEs in these processes is lagging behind because of an absence of tools amenable to their analysis. Here we provide the first 3C-based approach for addressing this question. 4Tran is important for learning how TEs such as MER41 influence transcription of host genes, and this in turn provides new insight into the mechanisms by which TEs contribute to the evolution of gene regulatory networks. Furthermore, 4Tran can be used to study the impact of TEs in disease settings such as cancer where the methylation status of the genome is altered and transposons become activated [75, 76].

## Conclusions

It is well established that TEs represent an important source of genetic variation [40]. These mobile elements were first identified in the late 1940’s by Barbara McClintock, who named them controlling elements for their ability to affect gene expression in maize. [37, 38]. However, it has only recently become apparent that TEs bound by transcription factors [33, 39] can have a major impact on gene regulation [40, 43]. Indeed, the spreading of multiple TF motifs in the genome by transposons is thought to be important in driving the evolution of gene regulatory networks. In addition, TEs have contributed to the organization of mammalian genomes by propagating binding sites, for the architectural protein, CTCF [33, 77]. However, it should be noted that not all TE integrations influence chromosome folding and gene expression as some do not harbor binding sites for regulatory factors and those that do, are often actively repressed

[19] and/or lose these sequences over time [44]. We developed 4Tran to characterize interactions between TEs and neighboring regions and to determine how they are physically organized in the nucleus. Here we show the usefulness of this approach in identifying contacts between transposons and the promoters of genes whose expression these elements have the capacity to control.

## Methods

### Mice

All mice used here were of wild-type inbred strains. Animal care was approved by Institutional Animal Care and Use Committee. Protocol number is 160704-02 (NYU School of Medicine). Young mice of less than 12 weeks were used in all experiments.

### Preparation of template for 4Tran-PCR

4C-Seq material was prepared from the following murine cells: embryonic stem cells, embryonic fibroblasts and splenic B cells. Starting material for each sample was 10 million cells. The 4C template material prepared from mouse embryonic stem cells was previously described [13]. These ES cells are from the ATCC cell line #CRL-1821 and were obtained from a 129/Ola mouse strain. Mouse embryonic fibroblasts were prepared from E13.5 embryos of the 129S1 strain. Embryos were isolated from timed-matings and dissociated using trypsin after removal of internal organs and decapitation. Cells were cultured for 2 passages before 4C material preparation. Resting mature splenic B cells were isolated as previously described [78] using magnetic beads for CD43 depletion from either C57/Bl6 Taconic mice (C57BL/6NTac) or from 129S6 mice also from Taconic (129S6/SvEvTac). Processing of 4C material was performed as described previously [58] using Dpn2 and Csp6I as enzymes for DNA digestion. Briefly, cells were fixed with formaldehyde and digested in situ with Dpn2. Following cell-lysis DNA fragments were diluted to favor proximity-mediated ligation. Concatemers of DNA fragment ligations were subsequently digested with Csp6I and religated upon dilution. 4C template was then amplified by thirty PCR cycles using 1µg of DNA divided by ten 50ul PCR reactions. Data is not shown here but we have also successfully tested *in situ* ligation by simply omitting the SDS treatment step following digestion with the primary restriction enzyme. Library amplification was done using Illumina single end adaptors, however the protocol also works with paired-end adaptors which are necessary for Illumina machines (Nextseq and Miseq for example). A list of samples generated in the study can be found in **Table S1** and a list of primers used for amplification can be found in**: Table S2.**

### Bait design and 4Tran-PCR

For each TE, 4C-Seq primers were designed to the consensus sequence obtained from Repbase. We then designed our algorithm to find all possible primer pairs based on the desired enzyme digestion. To be considered as a potential bait the following criteria are required: 300bp separating primary enzyme restriction sites and 150bp between primary and secondary enzyme restriction site. For these locations a 20-nucleotide primer is designed that includes the primary restriction enzyme site and the second primer is designed using the default primer 3 parameters of Primer3.

### 4Tran-PCR analysis

Single-end reads were mapped to a reduced genome of unique 25bp fragments adjacent to DpnII sites (mm10) using bowtie2 [79] as per the details described in 4C-ker [60]. Reads that map to the Encode blacklist regions were removed from downstream analysis [80]. To remove fragments that arise from self-ligation or incomplete digestion, only counts below a quantile of 99.9% were retained. For visualization, we generated counts in 100kb windows, overlapping by 25kb. In addition, to decrease the effect of PCR artifacts, counts in each window that are greater than the 75% quantile were reduced to that value. To define "observed baits" we used read counts obtained in 100kb non-overlapping windows. A z-score was then defined for each window based on the mean and standard-deviation across all windows in the genome. Finally, an FDR adjusted p-value was calculated based on fitting to a normal distribution and enriched windows were defined based on p-values smaller than 0.05. Windows separated by 100kb were merged. Only those windows identified in both replicates were called as observed baits. To determine the location of predicted baits we used the UCSC in-silico PCR tool. For that one of the primers for each bait was input as the reverse complement and the lowest stringency mismatch option (15 nucleotides) was chosen. To determine the regions interacting most frequently with specific TE integration events (**Figure 5**) we used the 4C-ker pipeline by performing cis analysis using a k of 10.

### Preparation of Capture-4Tran material

Murine B cell material was prepared as described in the section above. Human HeLa F2 cells were generous gifts from Cedric Feschotte and Edward Chuong. Cells were treated for 24 hours with 1000 U/ml of recombinant human IFNγ (cat# 11500-2, PBL assay science). Following treatment cells were fixed with Formaldehyde for 4Tran-Capture. To verify that cells responded to interferon treatment, RNA was collected and expression of the *IFI6* gene was measured by reverse transcriptase quantitative PCR. Primers can be found in **Table S2**.

### Capture-4Tran

Our approach was adapted from two published protocols. The Hi-C part of the protocol was performed mostly as described in [81]. Ligation of adaptors and Ilummina indices as well as DNA capture were done as detailed in [49]. Briefly, 10 million HeLa cells per replicate and condition were washed with PBS, trypsinized and spun down before resuspension in media containing 2% Formaldehyde. Fixation was done for 10 minutes at room temperature and stopped by adding Glycine. Nuclei were obtained after lysis on ice for 30 minutes with 10 mM Tris-HCl pH8, 10 mM NaCl, 0.2% Igepal CA-630 and 1 Roche Complete EDTA-free tablet. After 10 minutes treatment with 0.1% SDS at 37°C, nuclei were incubated with Triton 10% for 10 minutes also at 37°C. Nuclei were then suspended in 1X DpnII buffer (NEB) and chromatin digested overnight with 400U of DpnII (NEB) at 37°C. Following an additional 4-hour incubation with extra 400U of DpnII, digestion was stopped by 65°C incubation for 30 minutes. DNA ends were then filled in with dATP-Biotin using polymerase klenow for 60 minutes at 37°C. Ligation was performed *in-situ* overnight (with nuclei intact) using 2000U of T4 DNA ligase from NEB (M0202) at 16°C. Decrosslinking was performed overnight with Proteinase K at 65°C. Following two phenol chloroform extractions the efficiency of biotin fill in was verified using PCR and digestion with ClaI (Primers in **Table S2**). 40 μg of unligated biotin ends were removed by incubation with T4 DNA polymerase for 4 hours at 20°C. DNA fragments were then sonicated using an LE Covaris 220 instrument using 450 PIP, 30% duty factor, 200 cycles per burst for 60 seconds. Following sonication, DNA fragments were end-repaired using T4 DNA polymerase, T4 PNK and Klenow polymerase (all from NEB). DNA fragments were then bound to Streptavidin C1 beads and washed following manufacturer’s instructions and using 150 μl per 2.5 μg of sonicated DNA. A-tailing and adaptor ligation was then performed with DNA fragments bound to beads. A-tailing was done using Klenow polymerase without 3’ exonuclease activity. For ligation, adaptors from the NEBNext DNA library prep reagent set (E6040) were used and ligated overnight. Following ligation, enzyme USER treatment was used to open the stem loop. Hi-C library amplification was then amplified using 7 PCR cycles, with Phusion polymerase and PCR primers from NEBNext Multiplex Oligos for Illumina (E7335) and resulted in approximately 1 μg of Hi-C material. For the Hi-C sample, preparation was stopped here. For Capture, we used the Nimblegen SeqCap EZ hybridization and wash kit (05634261001), Nimblegen SeqCap EZ accessory kit v2 and Nimblegen SeqCap EZ HE-oligo kit. Briefly, 1 μg of Hi-C material was incubated at 42°C for 48 hours with 2.89 μmoles of a MER41 biotinylated DNA oligo according to Nimblegen’s instructions. Following recommended washes, beads were amplified using 8 cycles, with the Post-LM-PCR Oligos 1 & 2 oligos described in Nimblegen’s kit and Kappa 2X PCR master mix. This yielded approximately 700ng of DNA, which were used for a second round of capture as described above for 24 hours. Final amplification of libraries following second capture was done using 5 PCR cycles. In total 20 PCR cycles were used to amplify the libraries and approximately 100ng of material were prepared for sequencing using Illumina Nextseq paired-end 50 base pair reads.

### Hi-C and Capture 4Tran-capture analysis

Processing of Hi-C and Capture-4Tran reads was performed using Hicup [82]. Reads were mapped to the hg19 genome. Prediction of sites that hybridize to the MER41 probe was done using Blastn from ensemble (ensembl.org) with default parameters, without Repeatmasker filtering and allowing for up to 5000 hits. To detect fragments with high number of long-range interactions, each end of a mapped read-pair was first assigned to a restriction fragment in the genome using juicertools. Any read pairs that were within 100 fragments were removed. For the remaining interactions, the number of unique reads was counted and a quantile cut-off of 0.0005 was used to select the observed fragments with a high number of reads. This number was selected based on the percentage of Dpn2 fragments that Blastn predicts which are enriched by our oligonucleotide. Chicago [64] was used to identify interactions using the default parameters with a score cut-off of 7.5 on 2 replicates. The Dpn2 fragments with high number of reads were used as the baits (anchors) in Chicago and any fragment that did not overlap with a MER41 element was removed from downstream analysis (48/992). HiGlass (higlass.io/) after Cooler processing (github.com/mirnylab/cooler) was used to visualize Capture-4Tran results. Normalization of the Capture-4Tran data was performed using DESeq2 on read counts in 1kb bins overlapping by 500bp in a 1MB region around the bait. Normalized bedGraph files were used for visualization of data in IGV.

Differential analysis for the 4943 interactions was performed using DESeq2. The input for the analysis was the counts in the interacting region for each sample (2 replicated per condition) and adjusted p-value cut off of 0.05 was used to call significant interactions. UpSet plots were generated using the R package to calculate the overlap between features [83].

### Published Hi-C data analysis and Capture-4Tran analysis

Paired-end reads were mapped separately using bowtie 2.0.1. Downstream analysis was performed using Homer default parameters to obtain the count matrix and PCA scores to determine A and B compartment composition.

### Published transcriptional data analysis of Interferon treatment

To identify genes upregulated by interferon treatment in HeLa cells we used a microarray dataset [84] as this study contained three different replicates for each condition and was therefore more reliable. To identify upregulated genes we used the GEO2R tool from the NCBI GEO website (ncbi.nlm.nih.gov/geo) and used as criteria a log fold change of 1 and an adjusted p-value of 0.05. To identify all genes that are potentially expressed in HeLa cells we used RNA-seq data (SRR2992615-48Hr Control, SRR2992616-48Hr IFNγ, SRR2992619-72Hr Control, SRR2992620-72Hr IFNγ) [85]. Single-end RNA-Seq data was mapped to the hg19 genome using Tophat v2.1.1 (−-no-coverage-search –N 0 −-b2-very-sensitive). Htseq-counts was used to count the reads per gene using the hg19 Refseq annotation of genes. The counts were then normalized using DESeq2 for sequencing depth and further for gene length by dividing the normalized count by gene length and multiplying by 1e3.

## Ethics approval and consent to participate

Not applicable

## Acknowledgments

The authors thank all members of the Skok lab and Christian L Mueller for discussions as well as Helen Rowe and Jef Boeke for insights into transposon biology and Varun Narendra for 4C template used in the initial tests. The authors also thank the New York University School of Medicine Division of Laboratory Animal Resources (DLAR) for mouse breeding, the NYU High Performance Computing (HPC) (Shenglong Wang) for computing technical support, and the Genome Technology Center (GTC) core for sequencing efforts.

## Authors’ contributions

RR, PPR, RB and JAS conceived the study and designed the experiments. RR, PPR, VML and ES performed experiments. RR and PPR analyzed the data. ERM, EBC and CF helped with initial planning of experiments. RR, PPR and JAS wrote the manuscript. All authors read and approved the final manuscript.

## Supplementary tables

**Table S1** List of oligos used for 4Tran

**Table S2** List of 4Tran samples

**Table S3** List of Potential Regulatory Interactions.

**Fig. S1.**
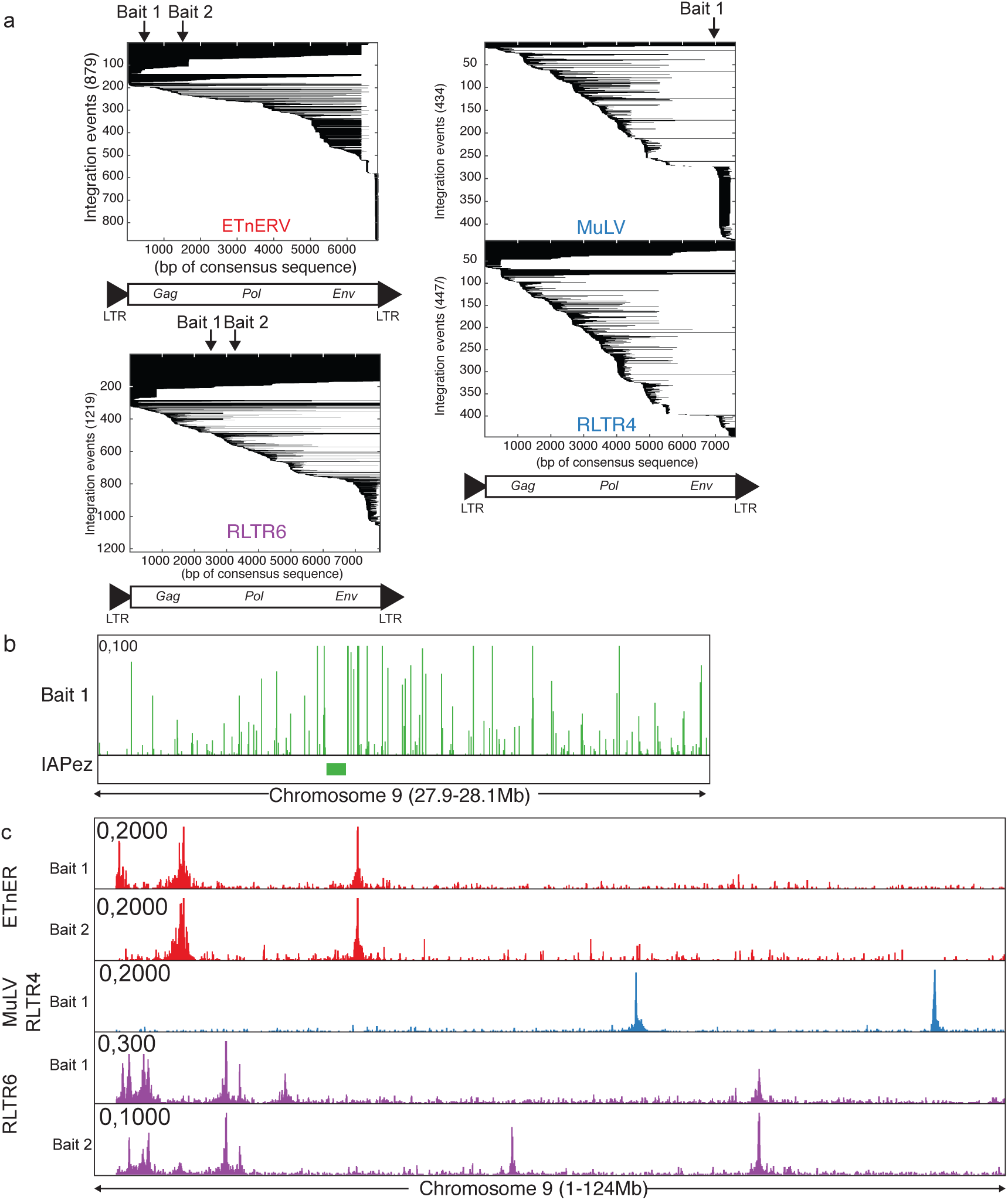
**a** Schematic representation of all integration events ERVs shown. Each line represents a different integration and the black shows which part of the consensus sequence (shown under the plot) is retained. Integration events are sorted by 5’ position on the consensus sequence and then by size of integration. Arrows represent the location tested for 4Tran-PCR baits. **b** Zoom in of a region on chromosome 9 showing raw 4Tran-PCR signal around an IAPEz integration (shown as a bar under the plot). **c** Raw 4Tran-PCR data for the baits tested in mES cells.

**Fig. S2.**
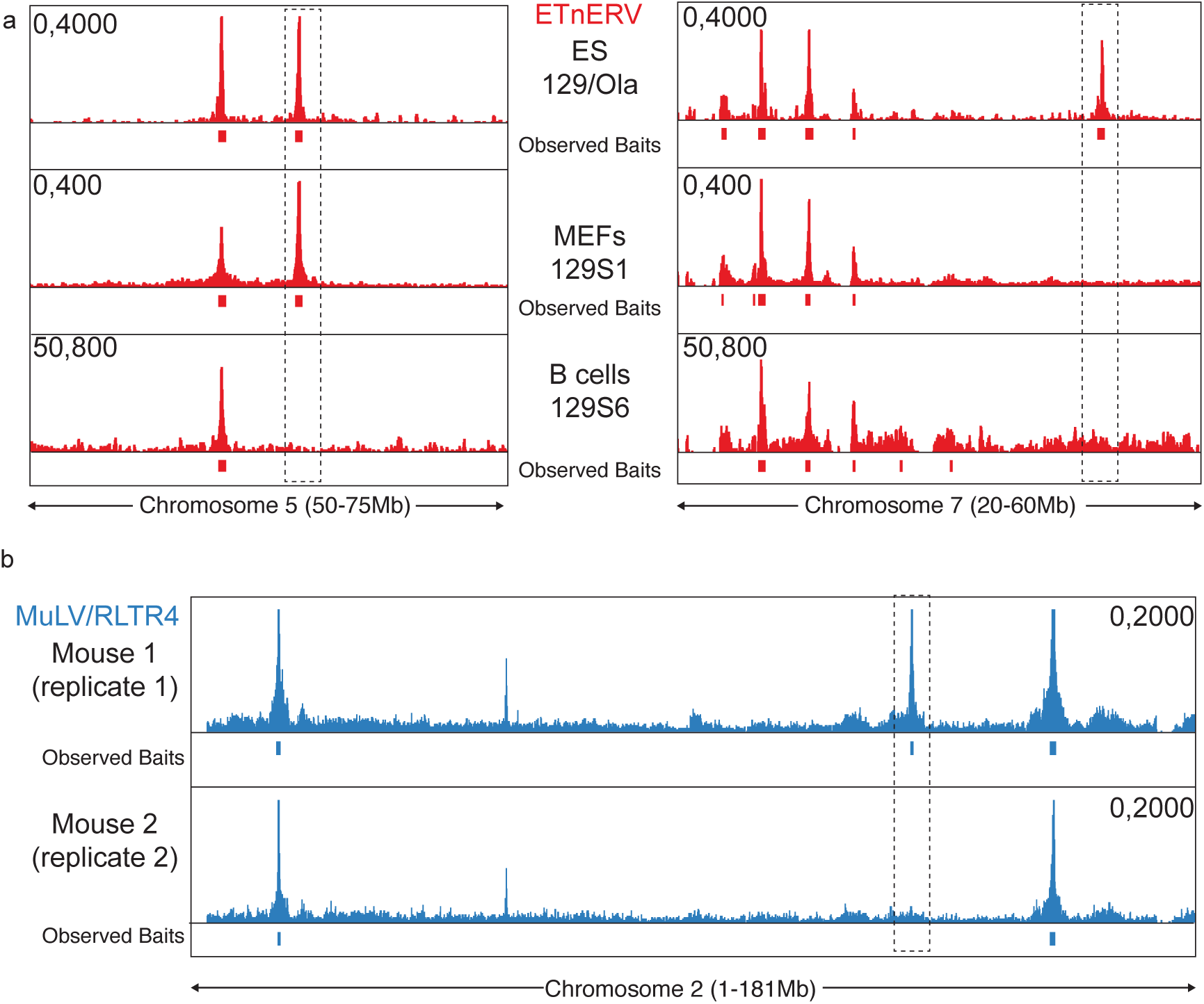
**a** Examples of shared and different integrations of ETnERV elements. The same primer pair was used to amplify signal from either mouse embryonic stem cells of the 129/Ola substrain, mouse embryonic fibroblast from the 129S1 substrain and splenic B cells of the 129S6 substrain. The locations of observed bait-like profiles is shown under the plots. **b** Example of a different integrations site detected in littermates.

**Fig. S3.**
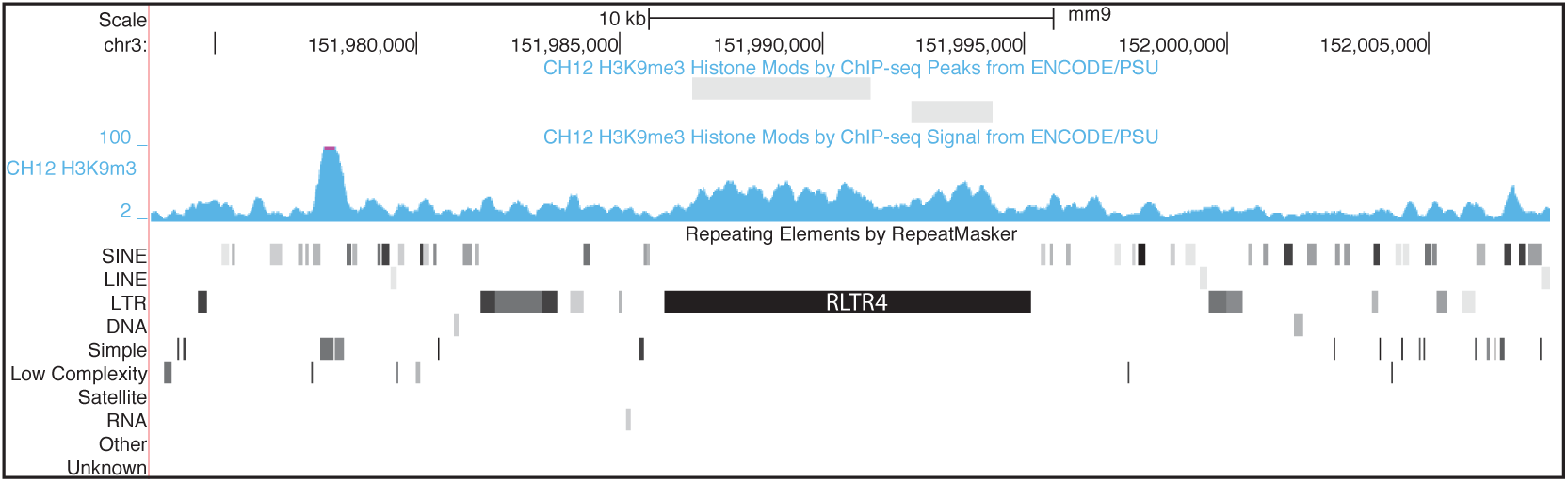
UCSC browser view of mouse chromosome 3 and of the ENCODE data for H3K9me Chip-Seq in the region surrounding the RLTR4 integration shown in Figure 4.

**Fig. S4.**
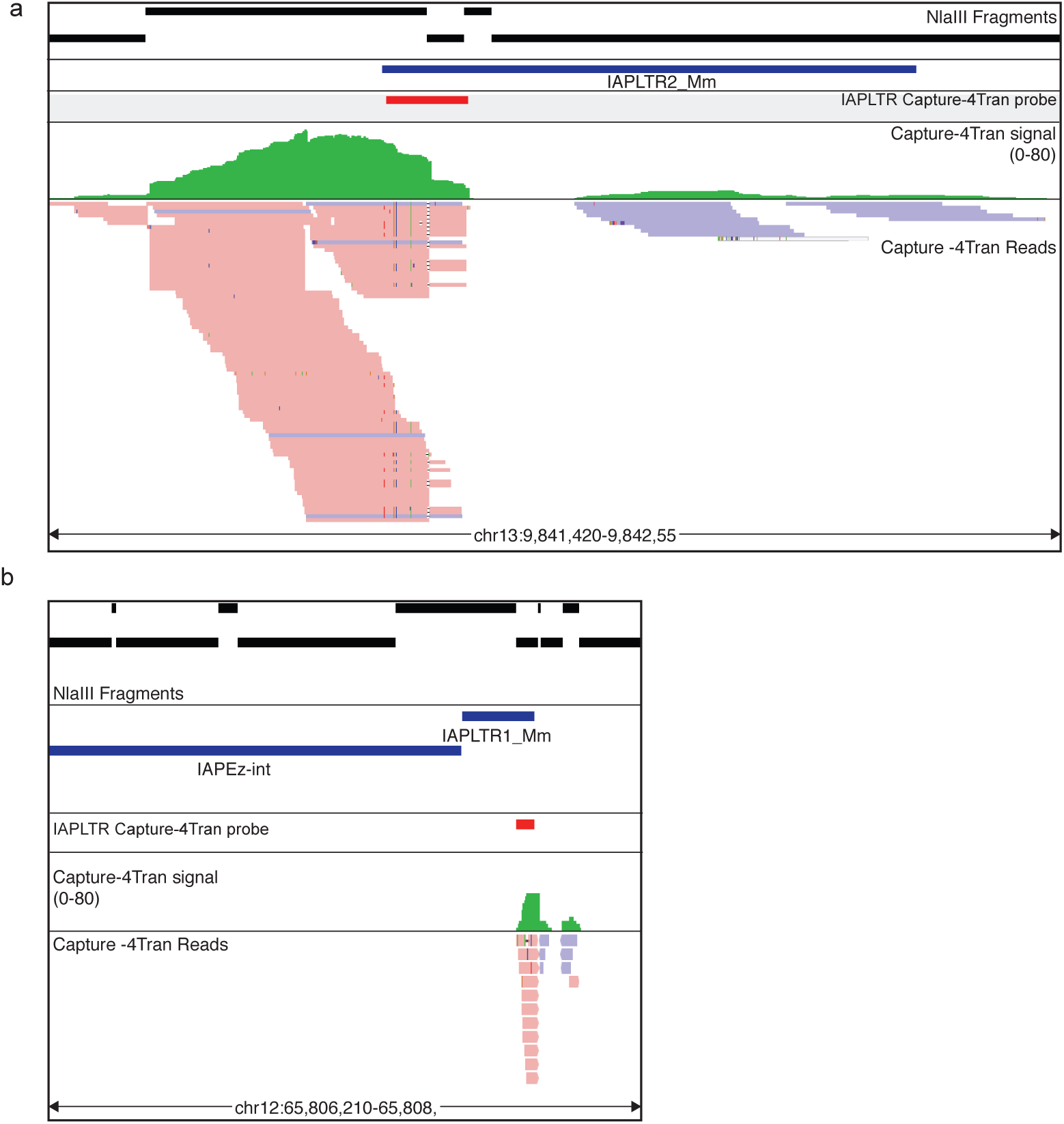
Browser view of Capture-4Tran using an IAPLTR probe on a solo LTR (**a**) and surrounding a full length IAPEz (**b**). Top track represents the DNA fragments generated by NlaIII digestion. The location of ERV elements followed by predicted and detected integration sites shown respectively as a peak profile or with actual location of reads is shown below.

**Fig. S5.**
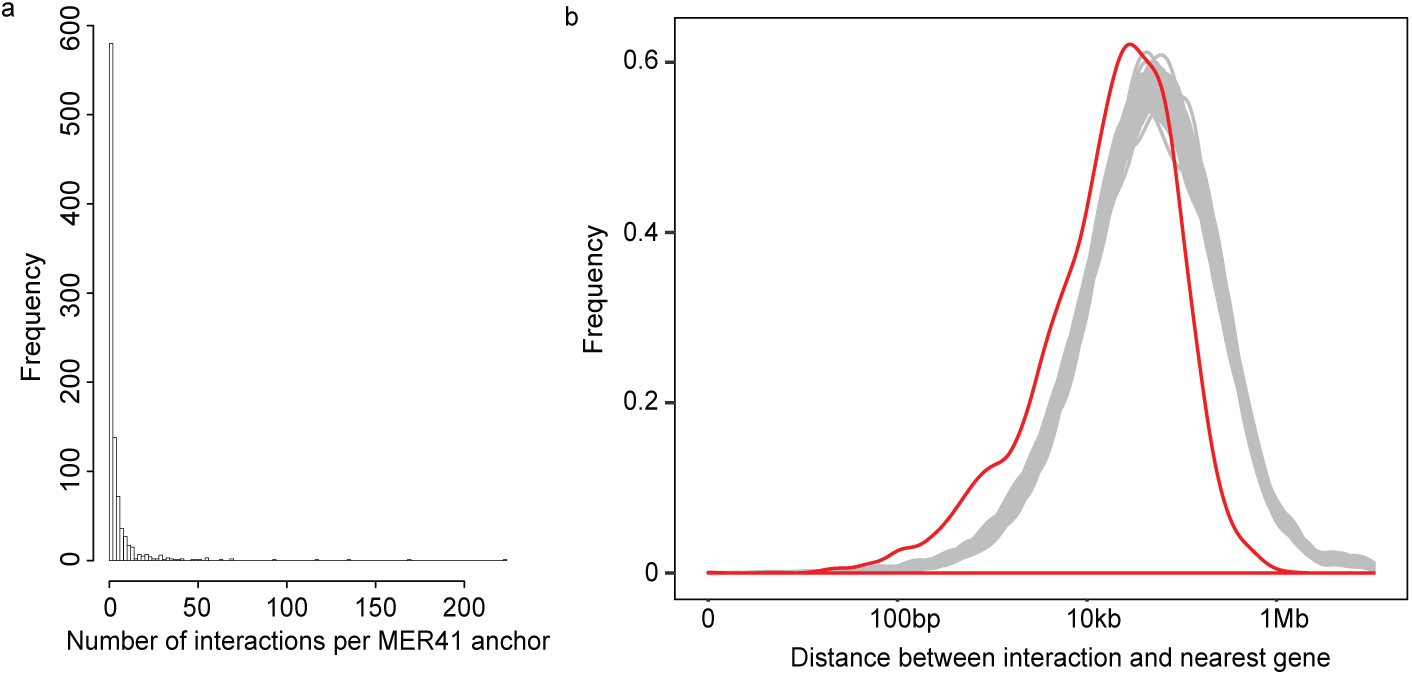
**a, b** Browser view of interactions captured from MER41 elements. Location and score of interactions detected using Chicago is shown as well as the location of the MER41 oligonucleotide used. STAT1 ChIP-Seq and RNA-Seq data were obtained from untreated and IFNγ treated HeLa cells. Data from two Capture-4Tran replicates for each condition was merged for visualization.

